# Patient-derived tumor organoids with p53 mutations, and not wild-type p53, are sensitive to synergistic combination PARP inhibitor treatment

**DOI:** 10.1101/2023.06.22.544406

**Authors:** Florencia P. Madorsky Rowdo, Gu Xiao, Galina F Khramtsova, John Nguyen, Olufunmilayo I Olopade, Rachel Martini, Brian Stonaker, Richard Boateng, Joseph K. Oppong, Ernest K. Adjei, Baffour Awuah, Ishmael Kyei, Frances S. Aitpillah, Michael O. Adinku, Kwasi Ankomah, Ernest B. Osei-Bonsu, Kofi K. Gyan, Nasser K. Altorki, Esther Cheng, Paula S. Ginter, Syed Hoda, Lisa Newman, Olivier Elemento, Melissa B. Davis, M. Laura Martin, Jill Bargonetti

## Abstract

Poly (ADP-ribose) polymerase inhibitors (PARPi) are used for patients with *BRCA1/2* mutations, but patients with other mutations may benefit from PARPi treatment. Another mutation that is present in more cancers than *BRCA1/2* is mutation to the *TP53* gene. In 2D breast cancer cell lines, mutant p53 (mtp53) proteins tightly associate with replicating DNA and Poly (ADP-ribose) polymerase (PARP) protein. Combination drug treatment with the alkylating agent temozolomide and the PARPi talazoparib kills mtp53 expressing 2D grown breast cancer cell lines. We evaluated the sensitivity to the combination of temozolomide plus PARPi talazoparib treatment to breast and lung cancer patient-derived tumor organoids (PDTOs). The combination of the two drugs was synergistic for a cytotoxic response in PDTOs with mtp53 but not for PDTOs with wtp53. The combination of talazoparib and temozolomide induced more DNA double-strand breaks in mtp53 expressing organoids than in wild-type p53 expressing organoids as shown by increased γ-H2AX protein expression. Moreover, breast cancer tissue microarrays (TMAs) showed a positive correlation between stable p53 and high PARP1 expression in sub-groups of breast cancers, which may indicate sub-classes of breast cancers sensitive to PARPi therapy. These results suggest that mtp53 could be a biomarker to predict response to the combination of PARPi talazoparib-temozolomide treatment.

## 1. Introduction

More than 50% of cancer patients present mutations in p53, and mutant p53 (mtp53) is correlated with advanced clinical stages and poor prognosis in multiple cancer types [1, 2]. However, diagnostic tools and treatment options targeted at the p53 pathway remain low and mutations in *TP53* are considered “undruggable”. *TP53* is the most frequently mutated gene in Triple Negative Breast Cancer (TNBC) (in about 80% of the patients) [3]. Moreover, 90% of small cell lung tumors are mtp53 [4]. The p53 protein is a tumor suppressor and DNA damage response (DDR) sensor which leads to cell cycle arrest, cellular senescence, DNA repair and apoptosis [5]. The most common type of *TP53* mutations are missense mutations and high occurrence “hot spot” mutant p53 (mtp53) proteins are R175, G245, R248, R249, R273, and R282. These missense mutations not only impair the tumor suppressive function of p53 but also have oncogenic gain-of-function (GOF) properties that promote cell proliferation, metastasis, genomic instability, metabolic reprogramming, and resistance to cancer therapy [6].

Unlike wild-type p53 (wtp53) protein, mtp53 is stable in cancer cells and mtp53 GOF activity requires elevated levels of mtp53 protein [7]. The GOF activity of mtp53 suggests that targeting the mtp53-dependent signaling network can be an effective approach for treatment. No mtp53-based therapeutics have been approved so far [8]. Our group investigates targeting the mtp53-PARP axis as a treatment paradigm and uses a cell penetrant tagged p53 tetramerization domain peptide called Cy5p53tet as a potential mtp53 diagnostic [9–13].

GOF mtp53 R273H up-regulates the chromatin association of PARP and minichromosome maintenance (MCM) complex MCM2-7 [12, 13] and interacts with replicating DNA [11]. This connects mtp53 to PARP-1 as a sensor of unligated Okazaki fragments during DNA replication in a potential cross-talk pathway [14]. We previously uncovered that inhibiting PARP activity induces cancer cell death selectively due to mtp53 mediated high level PARP association on DNA replication forks in cancer cell lines [11]. Therefore, we were interested in exploring the connection in a more clinically theranostic relevant model using patient-derived cells. Organoid models from diverse populations hold the promise to better understand and design precision medicine treatments for TNBCs, which are found to have high levels of genomic instability and cause high mortality in people of African descent [15–19].

Patient-derived tumor organoids (PDTOs) are 3D cellular models grown in culture *ex vivo* from the tumor tissues obtained from patients. PDTOs maintain the characteristics of the original tumor, as mutational signatures and tumor-clonal heterogeneity [20, 21]. They are valuable models for precision medicine approaches in drug screening and drug discovery. Several studies have shown a correlation between the patient and tumor organoid treatment response across several cancer types, showing that PDTOs have the capacity to replicate and predict treatment outcomes [20, 22, 23]. Representation of the global community in PDTOs is critical when evaluating cancers. As such, this study includes organoids collected both in the United States and in Ghana.

Recent evidence suggests that patients with wild-type *BRCA1/2* but presenting alternative HR deficiencies or mutations in genes involved in the DNA damage response can obtain clinical benefit from PARP inhibitor treatment alone or in combination with chemotherapy [24, 25]. Thus, the identification of biomarkers of response to PARPi remains a critical goal. Talazoparib is a potent PARPi, with both a strong catalytic inhibition of PARP and significantly more cytotoxicity by trapping PARP-DNA complexes than Olaparib [26]. PARPi combination with DNA damaging agents strategies can expand drug usage and overcome drug resistance [27].

The TCGA database and previous TMA data show high double positive PARP and mtp53 in basal-like breast cancer [11]. Herein our expanded molecular pathology studies from additional breast cancer tissue microarrays (TMAs) showed a positive correlation of high p53 and high PARP scoring in luminal A breast cancer subtypes. This further suggests that cancers can potentially be screened for mtp53-PARP dual biomarkers as a method to predict sensitivity to combination of PARPi talazoparib and the DNA damaging agent temozolomide. In order to test this hypothesis we used breast and lung cancer PDTOs to monitor PARPi response and revealed that the combination of PARPi talazoparib and the DNA damaging agent temozolomide induced a synergistic cytotoxicity in cancers with mtp53, but not wtp53, independent of *BRCA* mutations. The combination of talazoparib and temozolomide in a multiple concentration matrix uncovered synergistic cytotoxicity specifically in breast and lung PDTOs with mutations in the *TP53* gene derived from both breast cancer and lung cancer patients. Previously we designed a labeled peptide called Cy5p53tet to detect high levels of p53 (often correlated with missense mtp53 expression) in live cancer cells [28]. We detected higher uptake of the diagnostic peptide Cy5p53tet in organoids expressing mtp53 than in wtp53 expressing PDTOs. Thus, mtp53 and PARP expression may work as a dual prognostic biomarker in the absence of *BRCA1/2* mutations to guide the decision for combination treatment with talazoparib and temozolomide for cancer patients. As such the development of live cell imaging agents for such detection warrants further investigation.

## 2. Materials and Methods

### 2.1 Organoid establishment and culture

Patient-derived fresh tissue samples were collected with written informed patient consent with the approval of the Institutional Review Board (IRB #1305013903 and #1008011221) at Weill Cornell Medicine. ICSBCS002 and ICSBCS007 PDTO tissue samples were collected as part of the International Center for the Study of Breast Cancer Subtypes (ICSBCS) *Patient Cohort*. The ICSBCS is an international consortium of breast cancer clinicians and researchers with the broad goal of studying the heterogeneity of breast cancer across diverse population groups [29–31]. Institutional Review Board (IRB) approval has been obtained at all participating study locations, and in the present analysis with patients consented from the United States (Weill Cornell Medical College in New York City, NY) and Ghana (Komfo Anokye Teaching Hospital, Kumasi, Ghana) (IRB # 1807019405).

PDTO were developed as previously described [32] including some modifications. Fresh tissue samples were placed in CO_2_ independent medium (Gibco) with GlutaMAX (1×, Invitrogen), 100 U/ml Antibiotic-Antimycotic (Thermo Fisher Scientific) and Primocin 100 μg/ml (InvivoGen). Tissue samples were washed in culture media two times before being placed in a 10 cm petri dish for mechanical dissection. The dissected tissue was then enzymatically digested with collagenase IV media (DMEM (Gibco), 100 U/mL penicillin, 100 μg/mL streptomycin (Gibco), 250 U/mL collagenase IV (Life Technologies), 100μg/mL Primocin, and 10μmol/L Rock inhibitor Y-27632 (Selleck Chemical Inc.)) in a volume of at least 20 times the tissue volume and incubated on a shaker at 200 rpm at 37°C. Incubation time of the specimen was dependent on the amount of collected tissue and ranged from 20 to 60 min, until the majority of cell clusters were in suspension. After tissue digestion, Advanced DMEM/F12 media (Invitrogen) containing 1x GlutaMAX, 100 U/ml penicillin-streptomycin, and HEPES (10 mM, Gibco) (+++ media) was added to the suspension and the mixture was centrifuged at 300 rcf for 3 min. The pellet was then washed with +++ media and resuspended in breast or lung organoid-specific culture media (Supplementary table 1).

The final resuspended pellet was combined with Matrigel (Corning) in a 1:2 volume media:Matrigel, with 5 80 μl droplets pipetted onto each well of a six-well suspension culture plate (Greiner Bio-One). The plate was placed into a cell culture incubator at 37 °C and 5% CO_2_ for 30 min to polymerize the droplets before 3ml of organoid-specific culture media was added to each well. The culture was maintained with fresh media changed twice a week. Dense cultures with organoids ranging in size from 200 to 500 um were passaged every 10-30 days depending on the organoid line. During passaging, the organoid droplets were mixed with TrypLE Express (Gibco) and placed in a water bath at 37 °C for a maximum of 7 min. The resulting cell clusters and single cells were washed and re-plated, following the protocol listed above. PDTOs were cryopreserved in Recovery Cell Culture Freezing Medium (Gibco) in liquid nitrogen. Throughout organoid development and maintenance, cultures were screened for various Mycoplasma strains using the PCR Mycoplasma detection kit (ABM) and confirmed negative before being used for experimental assays. Whole exome sequencing (WES) and/or Oncomine targeted sequencing was performed on organoid cell pellets from passage 5 and matching tumors to confirm identity and concordance of mutational profile.

### 2.2. Drugs and peptides

Talazoparib was purchased from MedChemExpress and temozolomide was purchased from Selleckchem. Drugs were diluted and utilized according to manufacturers’ instructions. The working concentrations are indicated in each figure panel.

The Cy5p53Tet and Cy5Scramble were purchased from JPT peptide (Germany) at a purity >95%. The Cy5p53Tet is 37 amino acids long with an N-Terminal Cy5 fluorophore conjugation:

H-CysCy5-GRKKRRQRRRGEYFTLQIRGRERFEMFRELNEALELK-OH. The random scrambled peptide Cy5Scramble is 27 amino acids long designed by JPT peptide with an N-Terminal Cy5 fluorophore conjugation: H-Cys(Cy5)-LGEFELQKRQKRERLNGRLRERRAEFR-OH.

### 2.3. Drug dose-response assay

PDTOs were digested into a single cell suspension and cells were plated in a 384 well plate (Thermo Scientific, cat #164564) at a density of 1200 cells per well in 8ul droplets (1:2 media:matrigel). Plates were centrifuged briefly to ensure that the cells were at the bottom of the well and 15μl of media were added. Cells were incubated for 72h to allow organoid formation and the drugs were added at the indicated concentrations. After an additional 96h incubation, the viability readout was performed using CellTiterGlo®3D reagent (Promega) according to the manufacturer’s protocol. Luminescence was measured by the Biotek Synergy H4 plate reader. Non-linear regression analysis and IC_50_ determination were performed using GraphPad Prism 9 software.

For synergy analysis, the Synergyfinder web application 3.0 was used [33]. The synergy scores were calculated using the zero-interaction potency (ZIP) Synergy model. A synergy score lower than −10 indicates an antagonistic effect (green area), between −10 and +10 indicates an additive effect (white area), and higher than +10 indicates a synergistic effect (red area).

### 2.4. Cell fractionation

Cell fractionations were prepared using Chromatin Extraction Kit from Abcam (ab117152) according to the manufacturer’s protocol. Organoids were first lysed in Lysis Buffer (1X) containing proteinase and phosphatase inhibitors on ice for 10 min, vortexed vigorously for 10 sec and centrifuged at 2,300 g for 5 min. The supernatant was saved as the soluble fraction and extraction buffer was added to the pellet to resuspend it by pipetting up and down and incubating the sample on ice for 10 min, with occasional vortex mixing. The sample was resuspended and sonicated two times for 20 seconds/followed by 30 seconds of rest at 98% amplitude on ice to increase chromatin extraction. Then, the sample was centrifuged at 15,700 g at 4°C for 10 min. The supernatant was transferred to a new vial and Chromatin Buffer was added at a 1:1 ratio and saved as chromatin bound protein. The protein concentration was determined by Protein Bradford by NanoDrop One (Thermo Scientific).

### 2.5. Western Blot

Proteins were run on either 8%, 10% or 15% SDS-PAGE to separate samples followed by electro-transfer onto PVDF membrane (GE). The membrane was blocked with 5% non-fat milk (Biorad) in 1X Tris-Buffered Saline (TBS)/0.1% Tween-20 following incubation with primary antibody overnight at 4°C. The membrane was washed with 3 times of 1X TBS/0.1% Tween-20 and incubated with secondary antibody for 1 h at room temperature. The signal was detected by chemiluminescence with Super Signal Kit (Pierce) and autoradiography with Hyblot CL films (Denville Scientific) or ChemiDoc Imaging System (Bio-Rad).

### 2.6. Peptide cellular uptake by live cell imaging

Organoids were extracted from Matrigel using Cell Recovery Solution (Corning) following manufacturer’s protocol. Organoids were incubated with 500 nM Cy5p53Tet or Cy5Scramble at 37 °C for 4h. After incubation, the cells were washed three times with PBS at room temperature and co-stained with NucBlue live readyprobes reagent (Hoechst 33342) (Invitrogen). Cells were plated in 96 well CellCarrier Ultra imaging plates (Perkin Elmer) and imaging was performed using the Operetta High Content Imaging System (Perkin Elmer).

### 2.7. Whole cell protein extraction for Cy5p53Tet uptake analysis

Organoids were extracted from Matrigel using Cell Recovery Solution (Corning) following manufacturer’s protocol. Organoids were incubated with 500 nM Cy5p53Tet or Cy5Scramble at 37 °C for 4 h. After incubation, the cells were washed three times with PBS at room temperature. Organoids were harvested by centrifugation at 300 g for 3 min then whole-cell protein extraction from organoids were conducted as previously described [11].

### 2.8. Tissue microarray p53 and PARP1 immunohistochemistry

TMA were constructed from FFPE tumor samples and Immunohistochemical assays were performed as described previously [11]. Scoring was based on intensity and percentage of positive stained cells. The IHC immunoreactive score (IRS) was calculated based on percentage of positive cells and intensity of staining [34].

## 3. Results

### 3.1. Combination of PARPi talazoparib and temozolomide treatment yields synergistic cytotoxicity to mtp53 breast cancer PDTOs

Diagnostic and therapeutic interventions are critical for cancer treatments. We used breast cancer PDTOs to evaluate their sensitivity to PARPi talazoparib in combination with the DNA damaging agent temozolomide in the context of either wild-type or mutant *TP53*. This included PDTO lines with mtp53 R248W (ICSBCS002), mtp53 R181P (PM2438) and mtp53 R213stop (ICSBCS007) and two wtp53 breast PDTOs lines WCM2137_2 and WCM2384 (Table 1). We tested the sensitivity to a concentration gradient of talazoparib and temozolomide individually and found that ICSBCS007 was sensitive to talazoparib (IC_50_= 0.50 µM) while the rest of the PDTOs tested were resistant (IC_50_> 6.5 µM) (Figure 1A). For temozolomide, ICSBCS002 and ICSBCS007 presented some sensitivity (IC_50_=294.6 µM and 243.9 µM respectively) while the rest of the PDTOs were completely resistant (non-responsive) (Figure 1A). We then evaluated combination treatments using an inhibitor matrix to evaluate multiple combinatorial concentrations and calculated their synergy scores using the Synergyfinder3.0 web application (Figure 1B) [33].

**Figure 1.**
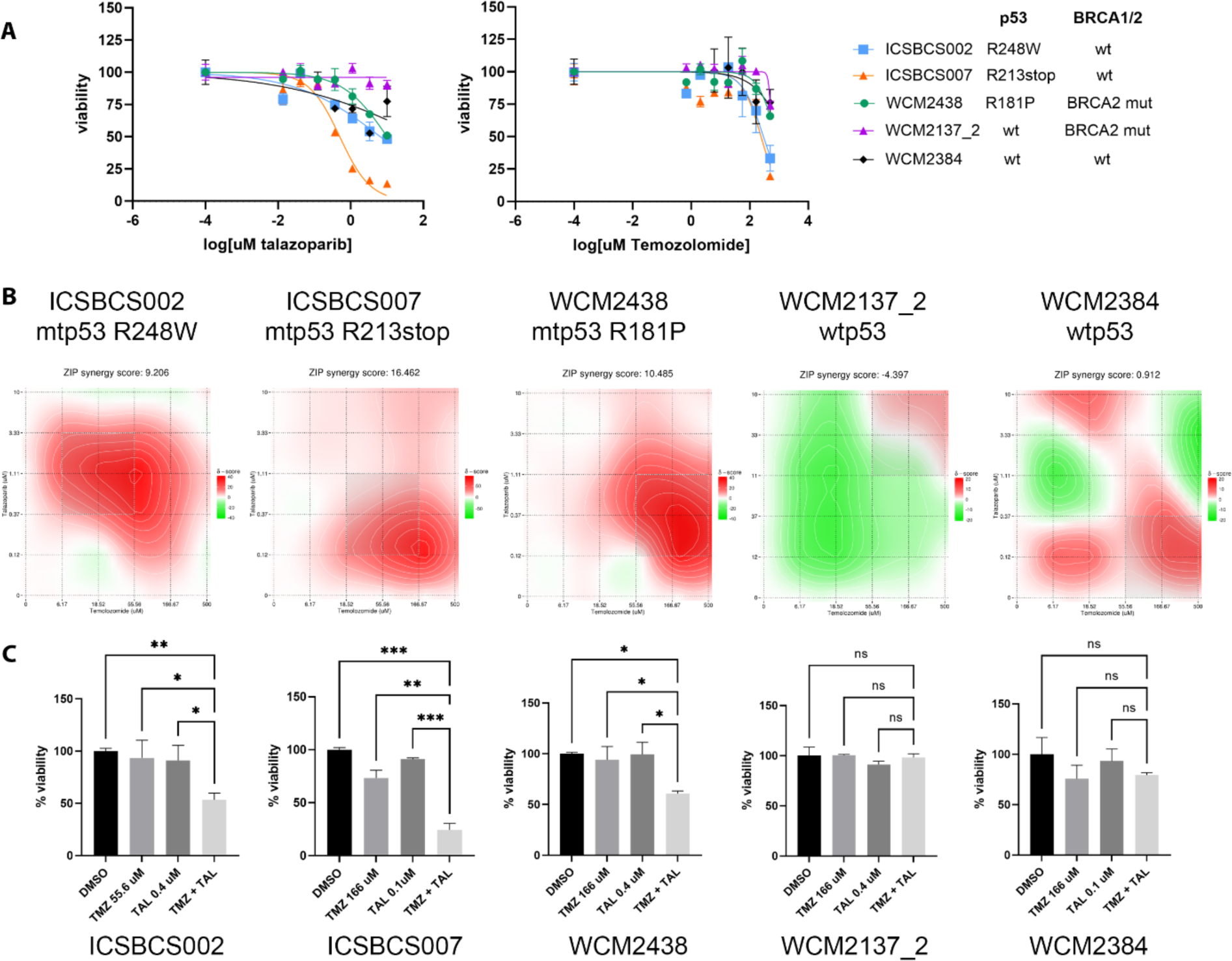
Combination PARPi talazoparib plus temozolomide treatment demonstrates synergistic cytotoxicity to mtp53 breast cancer PDTOs. (A) Drug response curves for breast cancer organoids to talazoparib and temozolomide. Cells were plated in 384 well plates and inhibitors were added after 3 days. After 4-day incubation viability was determined using Cell Titer Glo 3D reagent. (B) Synergy maps for talazoparib-temozolomide were calculated using the SynergyFinder web application with the ZIP synergy model (red indicates a synergistic effect, white an additive effect, and green an antagonistic effect). (C) Bar graphs showing cell viability at indicated concentrations.

**Table 1.**
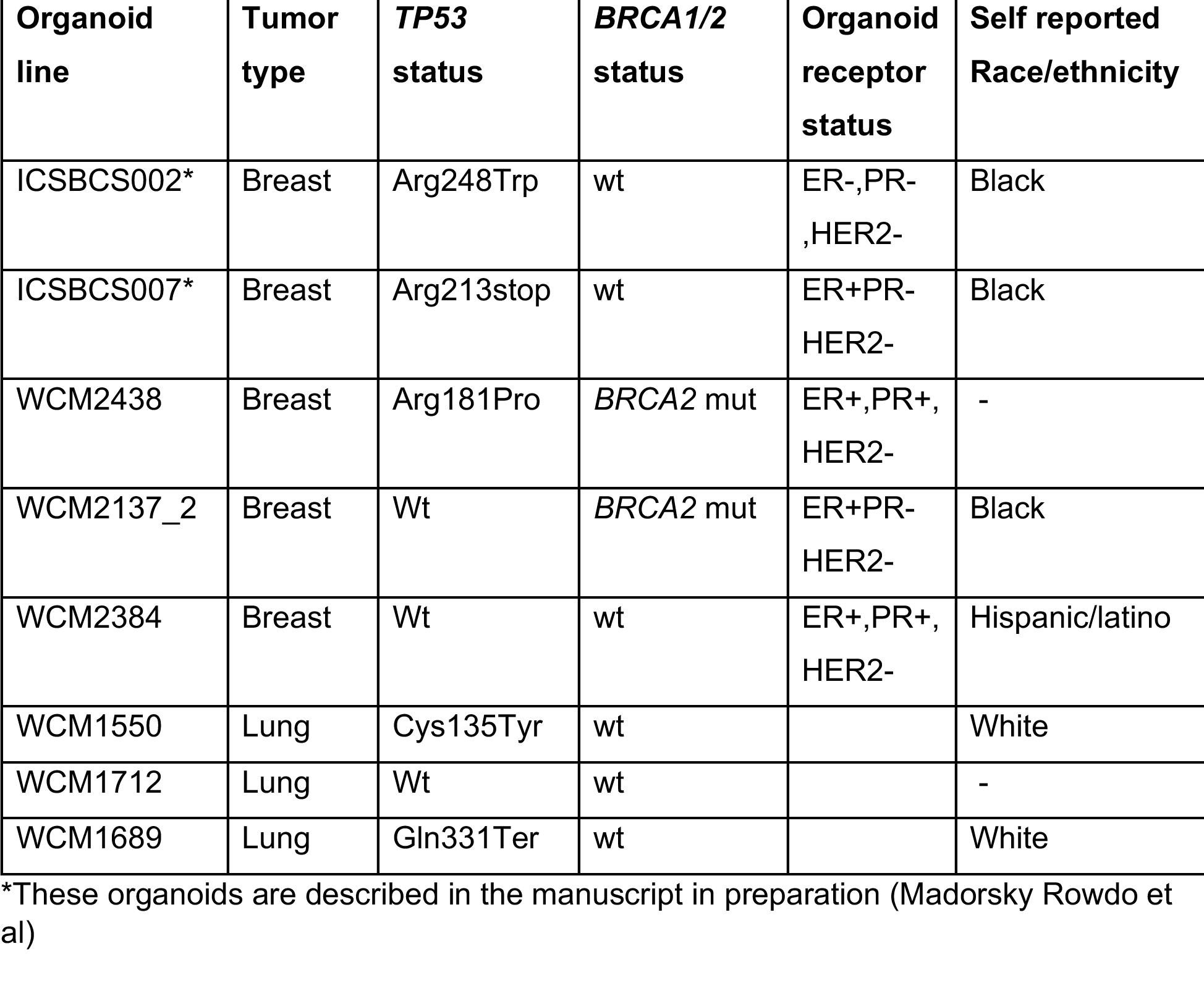
Patient-derived tumor organoid characteristics

Only the breast PDTO with mtp53 treated with the combination of temozolomide and PARPi talazoparib presented synergistic cytotoxicity (Figure1 B and C). A synergy score lower than −10 indicates an antagonistic effect (green area), between −10 and +10 indicates an additive effect (white area), and higher than +10 indicates a synergistic effect (red area). The two tested wtp53 breast organoid lines (WCM2137_2 and WCM2384) did not show synergistic sensitivity to the drug combination (Figure 1 B and C).The PDTO with mtp53 R248T (ICSBCS002) presented a global synergy score of 9.2, but for a selected range of concentrations the drug combinations presented a most synergistic area score of 20.35 (Figure 1 B and C). The PDTOs with mtp53 R213stop (ICSBCS007) and mtp53 R181P (WCM2438) presented scores greater than 10 (16.4 and 10.4 respectively) demonstrating mtp53 correlated synergy. This recapitulated our previous findings suggesting mtp53 is a good breast cancer biomarker for cytotoxicity of combination of PARPi and temozolomide treatment.

### 3.2. PARPi plus temozolomide synergistic treatment of mtp53 breast PDTO increases double strand breaks (DSBs) as indicated by chromatin bound γH2AX

To determine if the synergistic cytotoxicity induced by the treatment combination in mtp53 organoids, but not wtp53, correlated with increased DNA double-strand breaks (DSBs) we scored for phospho-H2AX at ser139 (γ-H2AX). DSBs activate the serine-threonine kinases that result in formation of increased γH2Ax foci and when cells have functional repair pathways, the damage is reduced [35]. We tested if breast cancer PDTOs treated with a combination of 55 µM temozolomide and 0.37 µM talazoparib for 24h (a dosage determined to be synergistic in Figure 1) resulted in more chromatin associated DSB marker γ-H2AX by evaluating chromatin and soluble protein respectively by western blot (Figures 2A and 2B). PDTOs with mtp53 but not wtp53 reproducibly presented increased levels of chromatin associated γ-H2AX (Figure 2A, compare lanes 4, 8, and 12). Treatments did not influence p53 protein levels which were highly stable for mtp53 R248W and barely detectable for both mtp53 R213stop and wtp53 (Figure 2, lanes 1-12 as labeled). Mtp53 R248W in ICSBCS002 was detected in both soluble and chromatin fractions but wtp53 in PDTO WCM2137_2 was only found in the chromatin fractions (compare lane 1-4 to lane 9-12). Talazoparib treatment of all PDTOs reduced total protein PARylation (Figure 2 see lanes 2, 4, 6, 8, 10, and 12). A significant increase of PARylated proteins were found in the wtp53 PDTO WCM2137_2 temozolomide treated chromatin fraction compared to mtp53 PDTOs (Figure 2A, lane 11), while temozolomide treated mtp53 PDTO soluble fractions presented high levels of PARylated proteins (Figure 2B, lanes 3 and 7). The highest PARP protein level was found in the “hot spot” mtp53 R248W ICSBCS002 in the chromatin fractions which supports our previous finding that highly stable GOF mtp53 interacts with PARP and promotes its tethering on the chromatin [11–13]. Taken together these data indicate that breast cancer PDTOs with mtp53 present increased sensitivity to combination of PARPi treatment that correlates with unrepaired DNA damage.

**Figure 2.**
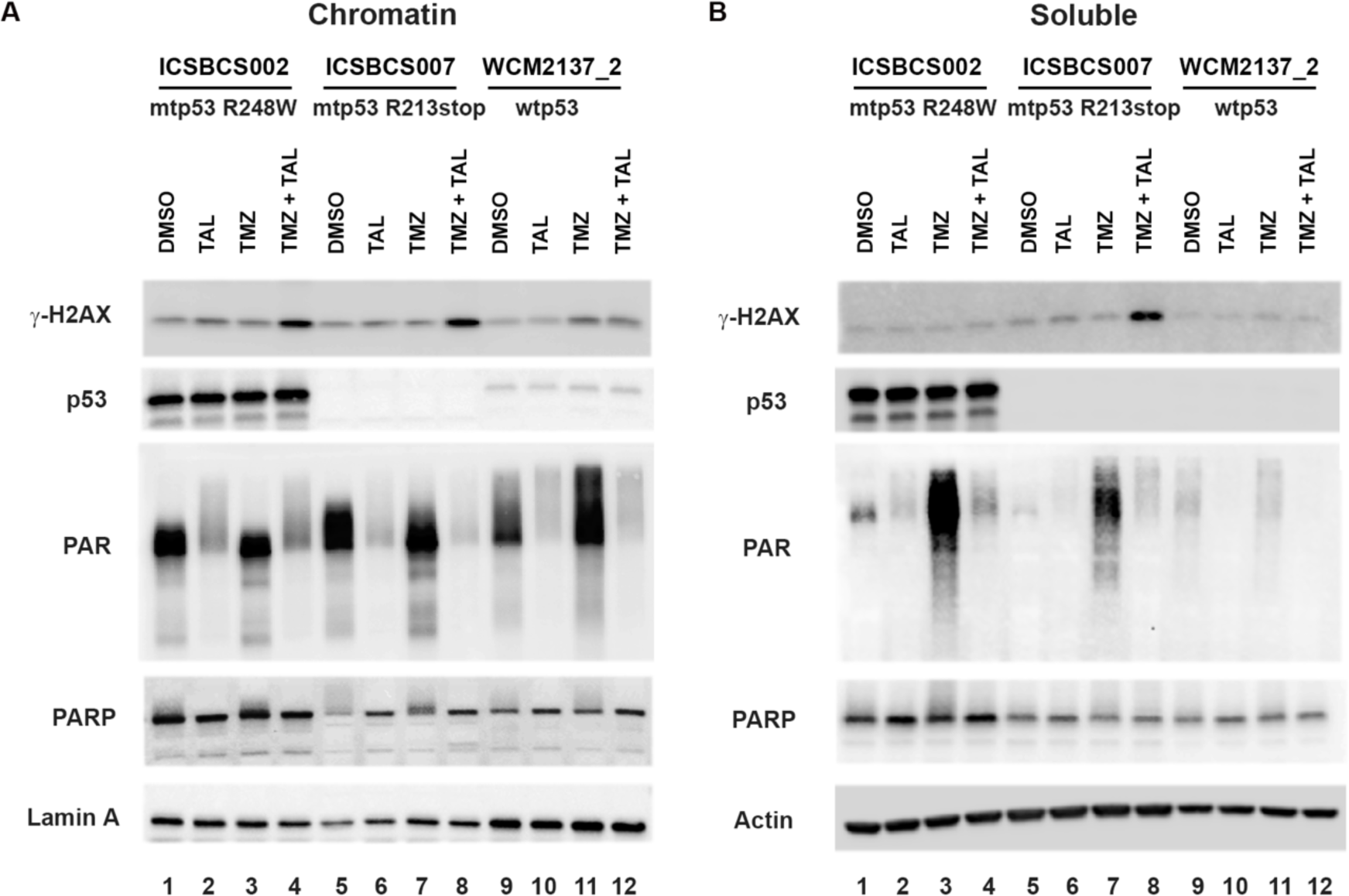
Synergistic PARPi talazoparib plus temozolomide treatment of mtp53 breast PDTO increases double strand breaks (DSBs) as indicated by chromatin bound γH2AX. Chromatin (A) and soluble (B) fractions were prepared from breast cancer PDTOs ICSBCS002 (mtp53 R248W), ICSBCS007 (mtp53 R213stop) and WCM2137_2 (wtp53) treated with either vehicle (DMSO), 0.4 µM talazoparib (TAL), 55 µM temozolomide (TMZ), or combination (TAL+TMZ) for 24 h. Protein levels of γ-H2AX, p53, PAR and PARP were determined by Western blot analysis.

### 3.3. PARPi plus temozolomide treatment results in synergistic cytotoxicity of mtp53 lung cancer PDTOs

We extended our studies to lung cancer PDTO with different status of p53 models to test the combination treatment of PARPi talazoparib and temozolomide. We asked if the mtp53-directed synergy of the combination of talazoparib and temozolomide could also be achieved in PDTOs of lung cancer. Lung cancers often have mutations in *TP53,* with 50–60% of non-small cell lung cancers and 90% of small cell lung tumors containing these oncogenic alterations [4, 36]. Lung cancer remains the leading overall cause of cancer mortality worldwide [37]. We compared non-small cell lung cancer PDTOs with mtp53 Q331stop (WCM1689), mtp53 C135Y (WCM1550), and wtp53 (WCM1712) (Table 1). The mtp53 Q331stop (WCM1689) presented sensitivity to single agent talazoparib (IC_50_=0.521 µM), while the two other PDTOs were resistant (WCM1550 IC_50_=4.134 µM and WCM1712 IC50=7.244 µM). All lung PDTOs were resistant to single agent temozolomide treatment (Figure 3A). However the matrix of combination dosing of temozolomide plus talazoparib presented mtp53-directed PDTO synergistic cytotoxicity (synergy scores >10) (Figures 3B and C). Taken together, these results indicate that PDTOs of both lung and breast cancer origin with mtp53 are more sensitive to the combination of talazoparib-temozolomide treatment than those cancers expressing wtp53. The wtp53 pathway responds rapidly to DNA damage and provokes cell cycle arrest and rapid DNA repair [38]. We therefore tested to see if the lung cancer PDTOs exposed to combination treatment showed correlated synergistic mtp53-associated increased DNA damage signal of γ-H2AX.

**Figure 3.**
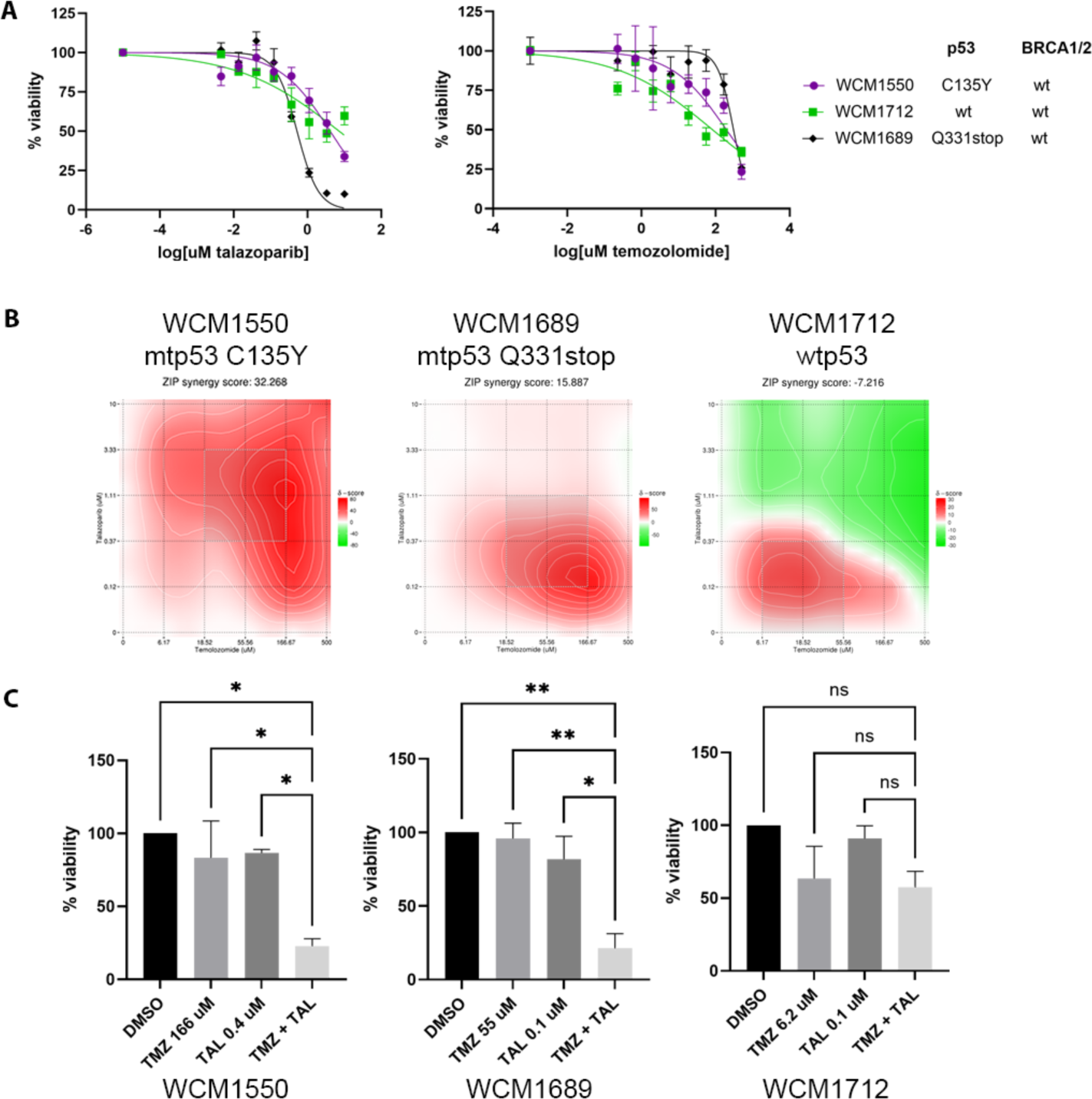
Combination PARPi talazoparib plus temozolomide treatment demonstrates synergistic cytotoxicity of mtp53 lung PDTO. (A) Drug response curves for lung cancer organoids to talazoparib and temozolomide. Cells were plated in 384 well plates and inhibitors were added after 3 days. After 4-day incubation viability was determined using Cell Titer Glo 3D reagent. (B) Synergy maps for talazoparib-temozolomide were calculated using SynergyFinder web application using the ZIP synergy model. (C) Bar graphs showing cell viability at indicated concentrations.

### 3.4. Synergistic PARPi plus temozolomide treatment of mtp53 lung PDTO increases DNA double strand breaks

To validate the molecular signaling of synergistic cytotoxicity of combination treatment we again analyzed γ-H2AX, p53, PARP and PARylated protein levels from the lung cancer PDTOs. We treated the lung cancer PDTOs with 166 µM of temozolomide and/or 0.4 µM of talazoparib for 24h as these concentrations showed synergy as demonstrated in Figure 3. Stable mtp53 Q331stop (WCM1689) and mtp53 C135Y (WCM1550) were detected in both the chromatin and soluble fractions and were undetectable in the wtp53 containing PDTO (WCM1712). Importantly, the combination of talazoparib and temozolomide induced more sustained DNA double-strand breaks in the mtp53 organoids WCM1550 compared to wtp53 WCM1712, as detected by increased chromatin bound γ-H2AX (Figure 4A). The chromatin bound γ-H2AX was increased following the combination treatments in comparison to single agent treatments and vehicle control for mtp53 PDTOs compared to the wtp53 PDTO (Figure 4A). In response to talazoparib treatment, the total protein PARylation (PAR) was reduced in all three PDTOs in both chromatin and soluble fractions (Figure 4A and 4B). Significantly, an increase of PARylated proteins were detected following temozolomide treatment (Figure 4A and 4B). Higher levels of PARylated proteins were found on chromatin in the wtp53 PDTO (WCM1712) compared to mtp53 PDTOs (WCM1689 and WCM1550). These results demonstrate that the combination of PARPi treatments results in sustained DNA damage in mtp53 PDTOs compared to wtp53 PDTOs pointing to this biomarker expression in both breast and lung cancers as an indicator of an enhanced cytotoxic effect due to impaired DNA repair pathways.

**Figure 4.**
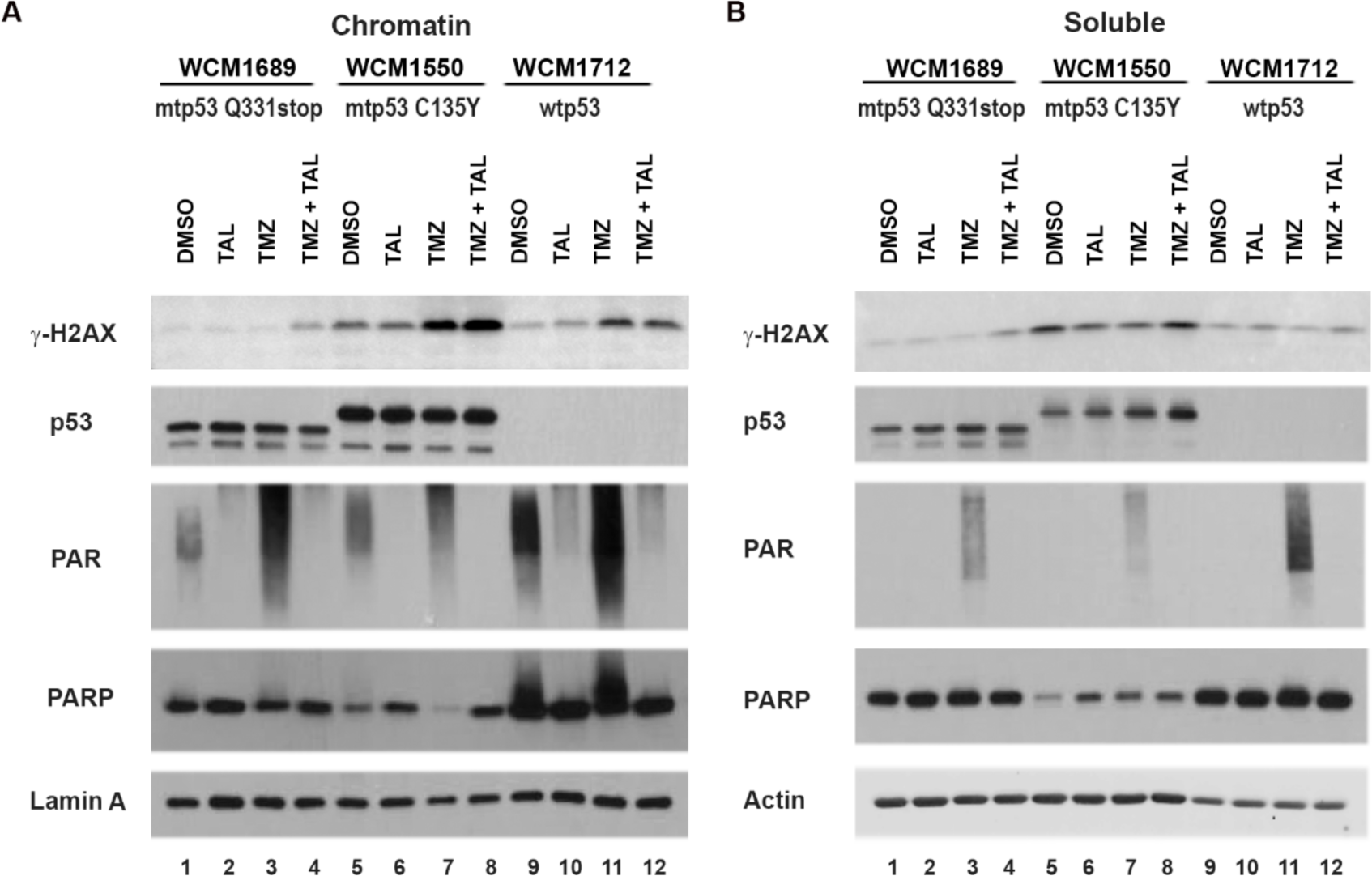
Synergistic PARPi talazoparib plus temozolomide treatment of mtp53 lung PDTO increases DNA double strand breaks (DSBs). Chromatin (A) and soluble (B) fractions were prepared from lung organoids WCM1689 (mtp53 Q331stop), WCM1550 (mtp53 C135Y) and WCM1712 (wtp53) treated with either vehicle (DMSO), or 0.4 µM talazoparib (TAL) or 166 µM temozolomide (TMZ), or combination (TAL+TMZ) for 24 h. Protein levels of γ-H2AX, p53, PAR and PARP were determined by Western blot analysis.

### 3.5. High mtp53 expression correlates with high PARP expression in TNBC and a sub-group of Luminal A breast cancers

TNBC lacks detectable Estrogen Receptor (ER), Progesterone Receptor (PR) expression and HER2 gene amplification [39], which causes very limited targeted therapies available for the treatment of patients with TNBCs. We previously found from TCGA data that p53 and PARP1 protein levels are significantly elevated in TNBC [11] and suggested that scoring mtp53 and PARP protein biomarkers may work as prognostic indicators for TNBC patients who may benefit from combination of talazoparib plus temozolomide treatment. We screened additional ethnically diverse patient tissue using breast tissue microarrays (TMAs) and scored for PARP and p53 protein expression using immunohistochemistry (IHC) (Figure 5 A-D). p53 IHC staining was performed on 161 samples from patients diagnosed with intrinsic subtypes of breast cancer, Basal-like, Luminal A, Luminal B and HER2+/ER-. The IHC score named immunoreactive score (IRS) was calculated based on percentage of positive cells and intensity of staining [16]. The immunoreactive score (IRS) is a product of multiplication between the percentage of positive cells score (0–4) and intensity of staining score (0– 3) (Figure 5D table and Representative images). A statistically significant higher IHC score of p53 was present in Basal-like breast cancer samples than luminal A or HER2+/ER-subtypes tumor tissues (Figure 5A). We compared the correlation between p53 and PARP levels of both p53 and PARP IHC scores from 138 breast cancer samples with validated p53 and PARP IHC in the same patient (Figure 5B). The Pearson correlation analysis showed a correlation coefficient of 0.3224 (P < 0.0001) demonstrating high p53 expression levels were positively correlated with high PARP expression levels in numerous samples. We asked whether there were ethnically diverse correlations of the p53 and PARP proteins in different subtypes of breast cancer among self-reported AA and white patients (Figure 5C). In both AA Basal-like (n=32) and AA Luminal A (n=37) tumor samples, the p53-high group was also high for PARP intensity. However, in the white Basal-like tumor samples (n=5), PARP intensity was not significantly different between the p53-high group and the p53-low group. In white Luminal A tumor samples (n=33), the p53-high group was also high for PARP intensity. Taken together, these data indicate that high mtp53 expression correlates with high PARP expression at least for Luminal A breast cancers in both self-reported racial groups (Figure 5). However, in this small sample set only the AA TNBC group showed a trend for a high p53 and high PARP (Figure 5C). These results reinforce the idea of the potential benefit of treatment combination of PARPi.

**Figure 5.**
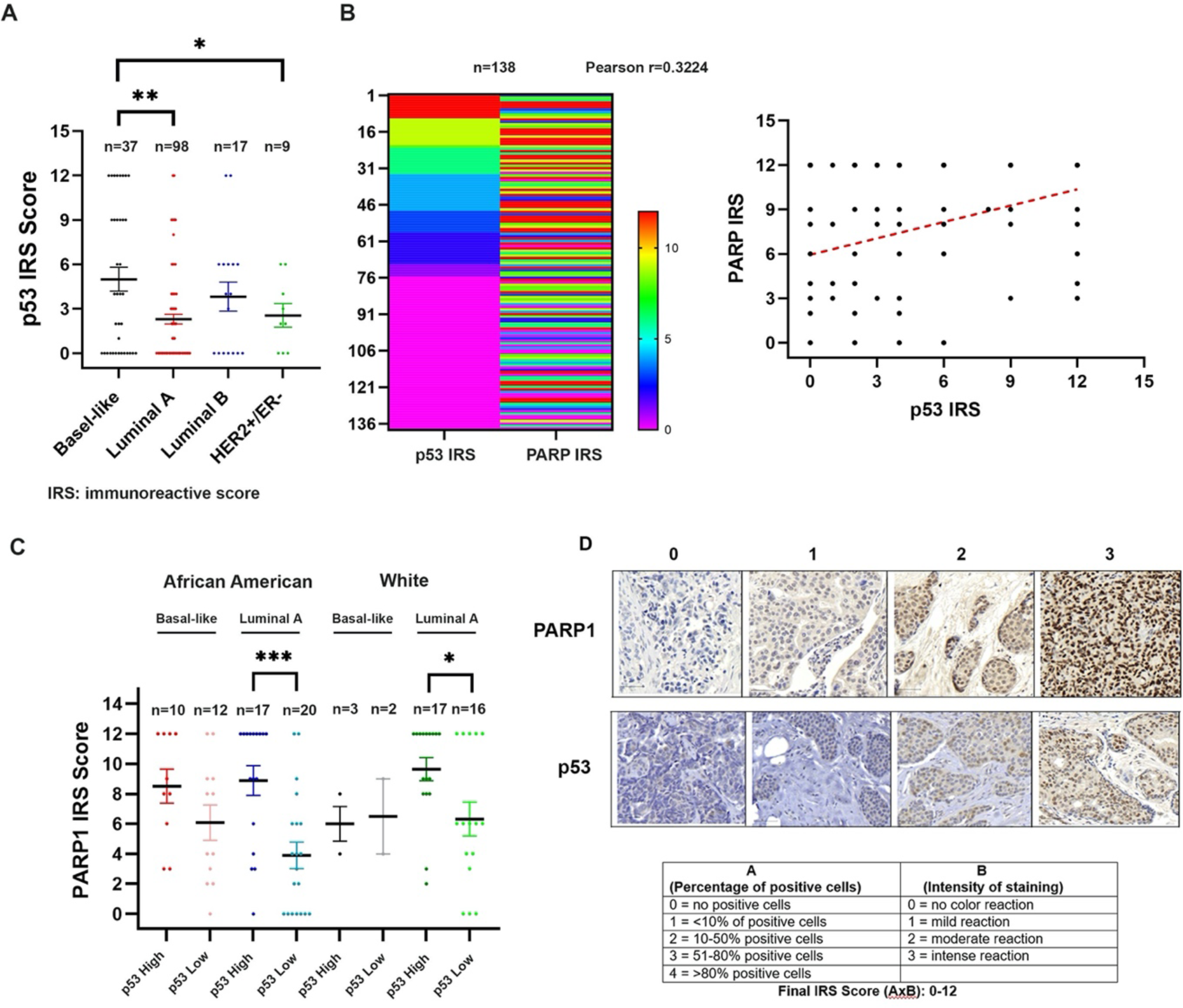
High mtp53 expression correlates with high PARP expression in TNBC of patients of African Ancestry. (A) Dot plot diagram showed IHC staining of p53 score (immunoreactive score: IRS) in tissue microarray (TMA) from Basal-like, Luminal A, Luminal B and Her2+/ER-subtype of breast cancer patients. The immunoreactive score (IRS) was calculated as shown in the table. (B) Correlation heatmap of p53 and PARP expression level (IRS) of 138 breast cancer patients. Each column of the data represents either p53 IRS or PARP IRS and each row represents one patient sample. Samples are ordered according to the p53 IRS level. The correlation of p53 and PARP was further analyzed using the Person correlation. y-axis presents the IRS of PARP and x-axis presents the IRS of p53. (C) Dot plot diagram showed IHC staining of PARP1 score in tissue microarray (TMA) from Basel-like and Luminal A subtype of breast cancer grouped by p53 level and ethic group (AA or white). (D) Representative images at 100x magnification of PARP and p53 staining. Intensity categories: no staining (0), mild staining (1), moderate staining (2) and intense staining (3). Scale bar: 50 um. The immunoreactive score (IRS) is a product of multiplication between percentage of positive cells score (0–4) and intensity of staining score (0–3). *P < 0.05, **P < 0.01, ***P <0.001.

### 3.6. Detecting mtp53 expression by specific uptake of Cy5p53Tet in breast cancer organoids to determine potential treatment response with temozolomide and talazoparib

Cancer care requires fine-tuned diagnostics and targeted therapeutics. As such, utilizing the most often mutated gene in cancers as a diagnostic and therapeutic target would change the nature of diagnosis and treatment for both lung and breast cancers. Elevated levels of mtp53 are found in tumor tissue while healthy tissues with normal functional p53 have undetectable levels of the p53 protein [6]. We designed a nuclear penetrant peptide to detect high levels of p53 protein by a fluorophore-linked peptide which we named Cy5p53Tet [28]. The peptide includes a nucleus-penetrating (HIV-TAT) amino acid sequence linked to the p53 tetramerization-domain (TD) structure that facilitates selectively targeting high levels of mtp53. *In vivo* analysis demonstrated that Cy5p53Tet specifically detects mtp53 expressing human xenograft tumors [28]. Herein we asked if the diagnostic Cy5p53Tet peptide could penetrate specifically into the PDTOs we studied for synergistic combination of PARPi and temozolomide. We performed live cell imaging to compare Cy5p53Tet to a scrambled Cy5-labeled peptide (Cy5Scramble) to stain the PDTO with mtp53 Arg213stop p53 expression (ICSBCS007) and the TNBC organoid with missense mtp53 Arg248Trp (ICSBCS002). The PDTOs were incubated with 500 nM Cy5p53Tet or the random scrambled Cy5Scramble peptide and co-stained with Hoechst for nuclear DNA staining (Figure 6A). The intensity of the Cy5p53Tet signal was significantly higher in the missense mtp53 Arg248Trp PDTO thus correlating with the mtp53 high stability (Figure 6B and see Figure 2). Quantification of the uptake of Cy5p53Tet and Cy5Scramble via Harmony (Perkin Elmer) analysis demonstrated a 1.7-fold higher uptake of Cy5p53Tet in missense mtp53 Arg248Trp ICSBCS002 compared to mtp53 Arg213stop ICSBCS007 PDTO. Both ICSBCS002 and ICSBCS007 organoids had 10-fold higher uptake of Cy5p53Tet than uptake of the non-specific Cy5Scramble peptides validating the diagnostic peptide specificity (Figure 6A&B). We further assessed the accumulation of Cy5p53Tet versus the Cy5Scramble in PDTOs by extracting proteins from ICSBCS002 and ICSBCS007 after incubation with Cy5p53Tet or Cy5Scramble and then quantitating the associated peptide level. Higher Cy5p53Tet was detected in ICSBCS002 compared to ICSBCS007 and only low levels of Cy5Scramble were detected. The difference of Cy5p53Tet uptake between ICSBCS002 and ICSBCS007 PDTOs was correlated to the level of p53 expression by Western blot (Figure 6C, compare lane 1-2 to lane 3-4). The uptake of Cy5p53Tet signal in ICSBCS007 suggested that Cy5p53Tet also interacted with other members of the p53 family, p63 and p73 that have high sequence and structure similarity in TD domain [40]. The TD domains of p53, p63 and p73 are dimers of dimers and the basic module consists of an antiparallel β-sheet formed by two monomers that is stabilized by two α-helices in an antiparallel orientation [40]. The protein expression levels of p63 and p73 were analyzed in ICSBCS002 and ICSBCS007 PDTOs (Figure 6D). Higher p63 and p73 protein expression were detected in ICSBCS007 PDTO compared to ICSBCS002. The higher protein expression levels of p63 and p73 in ICSBCS007 suggested that the detection of Cy5p53Tet could result from peptide binding also binding to p63 and p73. The increased uptake of Cy5p53Tet, and low uptake of scrambled peptide, was observed in lung PDTOs expressing mtp53 protein (Figure 6E). Further investigation of the protein interactions for Cy5p53Tet diagnostic are needed but our studies showed Cy5p53Tet can be used as a PDTO staining reagent for detection of stable high expression of mtp53 protein. Furthermore, Cy5p53Tet provided an image-based nondestructive approach to detect PDTOs based on mtp53 expression and might be able to predict the dynamic response of PDTOs to the combination treatment of temozolomide and talazoparib.

**Figure 6.**
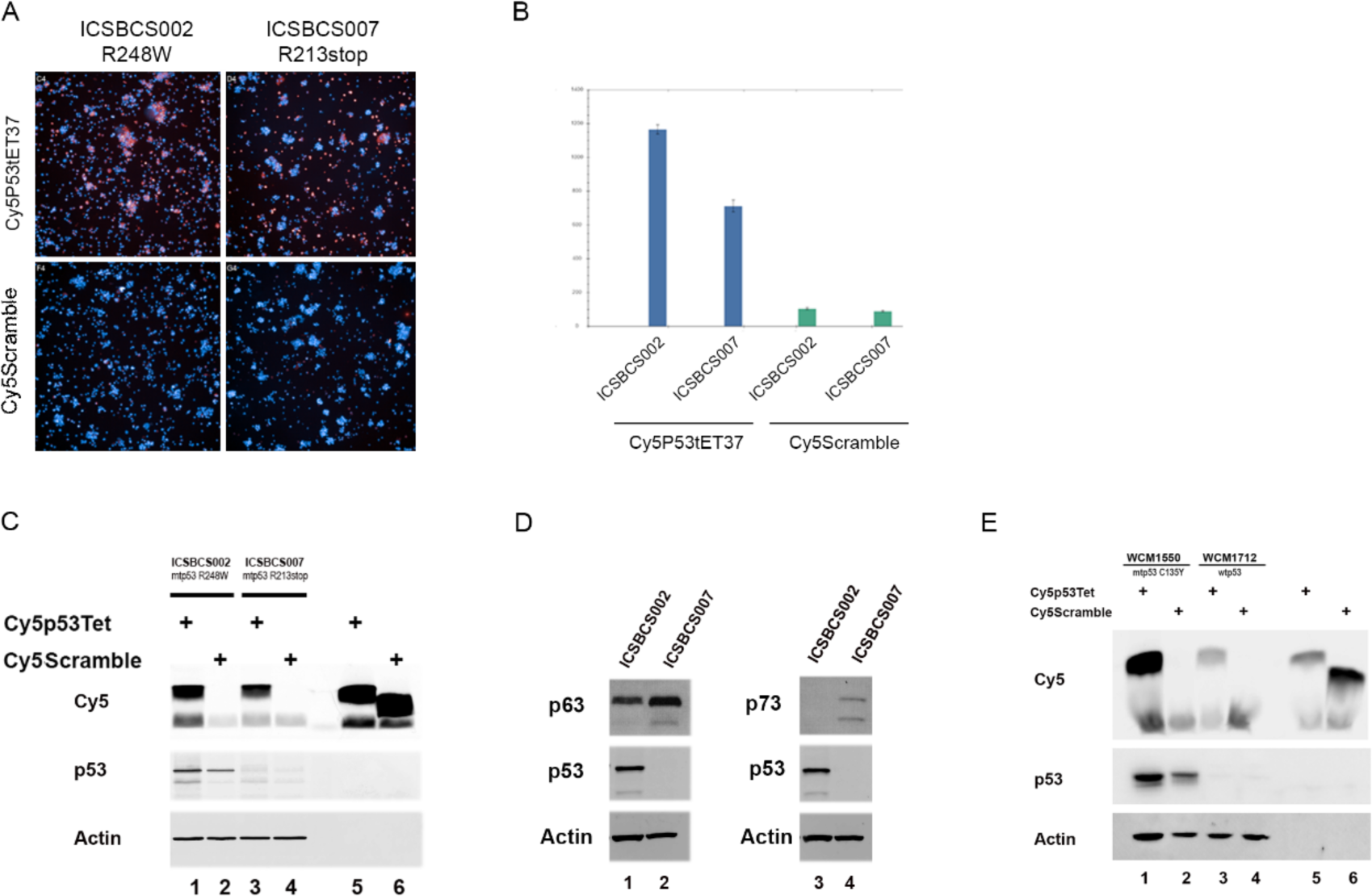
Specific uptake of Cy5p53Tet in high levels of mtp53 breast cancer organoids. (A) Representative images from high-content imaging analysis of mtp53 PDTO ICSBCS002 (mtp53 R248W) and ICSBCS007 (mtp53 R213 stop) treated with either 500 nM Cy5p53Tet or Cy5scramble peptides for 4 h. Hoechst staining was used to stain the nuclei (Cy5: red; Hoechst: blue). (B) Quantification of Cy5p53Tet and Cy5scramble uptake in ICSBCS002 and ICSBCS007 PDTOs via Operetta High Content Imaging System. (C) Whole cell extracts were prepared from breast cancer PDTO ICSBCS002 and ICSBCS007 treated with either 500 nM Cy5p53Tet or Cy5scramble peptides for 4h. 1 ug of cell extracts was run on 15% polyacrylamide gel and uptake of Cy5p53Tet or Cy5Scramble was examined by scanning the gel using Typhoon FLA 7000 biomolecular imager. Protein levels of p53 and Actin were determined by Western blot analysis. (D) Whole cell extracts were prepared from breast cancer PDTO ICSBCS002 and ICSBCS007. 25 ug of cell extracts were run on 10% polyacrylamide gel and protein levels of p63 and p73 were determined by Western blot analysis. (E) Whole cell extracts were prepared from lung cancer PDTO WCM1550 (mtp53 C135Y) and WCM1712 (wtp53) treated with either 500 nM Cy5p53Tet or Cy5scramble peptides for 3 h. 1 ug of cell extracts was run on 15% polyacrylamide gel and uptake of Cy5p53Tet or Cy5Scramble was examined by scanning the gel using Typhoon FLA 7000 biomolecular imager. Protein levels of p53 and Actin were determined by Western blot analysis.

## 4. Discussion

For over three decades (since p53 was deemed molecule of the year back in 1993) research has explored ways to solve the challenge of targeting cancers with mutations in the p53 pathway. With the advent of PDTO technologies the possibilities are here for faster drug development and precision medicine diagnostics due to physiological maintenance of cell-cell and cell-matrix interactions in these patient tissue models [41]. Herein we demonstrated on lung and breast PDTOs a synergistic drug combination that specifically targets mtp53 PDTOs. Combination drug synergism was achieved when treating mtp53 containing, but not wtp53 expressing, breast and lung PDTOs. Mechanistic insights indicated that synergy resulted from double-strand breaks that were not able to be repaired, and subsequently led to cell death. Importantly we found that the combination drug synergism occurred in both breast and lung PDTOs carrying wild type *BRCA1/2*. This demonstrated that using mtp53 as a significant biomarker for PARPi therapy independent of *BRCA* status is possible. Additionally scoring breast cancer patient tissue for the high expression of the p53 biomarker (thus detecting mtp53) and high PARP suggested dual biomarker screening as an option for the advancement of determining cancers that can respond to PARPi targeted therapy for beneficial clinical interventions.

The development of chemotherapy resistance during the progression of metastatic breast cancer is a profound problem that requires radical solutions [42]. Leveraging DNA damage stress pathways in breast cancers by inhibiting PARP are showing great promise [43, 44]. The data presented herein demonstrates the importance to determine how mtp53 interacts with DNA damage tolerance factors to provide strategies that target the replication stress responses for optimal cancer treatments [43]. Future studies utilizing PDTO models combined with comprehensive multiomics data: genetic mutation, gene expression, microRNA expression, copy number variation (CNV), methylation, and proteomics datasets generated from the same patients will provide opportunities to determine the pathways that provoke the synergy of combination PARPi therapy. While these PDTO models generated reproducible high quality PARPi sensitivity data potentially predictive of responses to treatment at the level of the individual cancer patient, limitations and optimization needs to be addressed before clinical implementation occurs. The absence of the tumor microenvironment (TME) interactions, and lack of vascular networks limits the potential to determine how the immunological response will impact outcomes [15]. Our current models are deprived of stromal and immune TME, therefore, TME effect to combination treatment must be incorporated into future studies. Co-cultures with cancer associated fibroblasts and immune cells will allow better interrogation of the intercellular interactions. Furthermore, a microfluidic device, Organoid-On-A-Chip models allow for vascularized organoid technology to recreate more complex *in vivo* dynamics of the extracellular environment including vascular, neuronal, and immune system [45]. A biobank of pre- and post-treatment PDTOs, which can be further used for drug sensitivity or resistance mechanism studies must be developed. This study warrants further implementation of combination of PARPi and temozolomide in high-throughput drug screenings using PDTOs. In perspective, we envision that the findings from our PDTOs will aid therapeutic decisions used by clinical practices for treatment of cancer patients with mtp53.

## Acknowledgements

This work was supported by The Breast Cancer Research Foundation BCRF-20–011 and BCRF-21–011 to J. Bargonetti, and by National Cancer Institute of the National Institutes of Health under Award Number R01CA239603 to J. Bargonetti. Research reported in this publication was also supported by the National Center for Advancing Translational Sciences of the NIH under Award Number UL1TR002384 and National Cancer Institute of the National Institutes of Health under Award Number T32CA203702. The content is solely the responsibility of the authors and does not necessarily represent the official views of the National Institutes of Health.

