## Supplementary table 1 for "Patient-derived tumor organoids with p53 mutations, and not wild-type p53, are sensitive to synergistic combination PARP inhibitor treatment"

Supplementary table 1. PDTO media composition.

| Media component |  | Final concentration |
| --- | --- | --- |
| Advanced DMEM/F12 | Thermo Fisher Scientific 12634028 |  |
| Glutamax | Life Technologies 35050079 | 1% |
| HEPES | Thermo Fisher Scientific 15630-080 | 1% |
| Penicillin/Streptomycin | Thermo Fisher Scientific 15140163 | 100U/ml |
| B27 supplement | Life Technologies 17504-044 | 1X |
| Nicotinamide | Sigma-Aldrich N0636-100G | 10mM |
| N-Acety-l-cysteine | Sigma-Aldrich A9165-5G | 1.25 mM |
| Primocin | Invivogen ant-pm-1 | 1X |
| recombinant human FGF-basic | Peprotech 100-18B | 1 ng/mL |
| recombinant human FGF10 | Peprotech 100-26 | 20ng/ml |
| PGE2 | R&D Systems 2296/10 | 1μM |
| SB202190 | Sigma-Aldrich S7067 | 10μM |
| mouse recombinant EGF | Thermo Fisher Scientific PMG8043 | 50ng/mL |
| Y-27632 | VWR S1049-50MG | 10μM |
| A-83-01 | VWR 10188-672 | 500nM |
| Heregulin beta-1 | PeproTech 100-03 | 10 ng/mL (lung) or 200ng/ml (breast) |
| Noggin conditioned media |  | 10% |
| R-spondin conditioned media. |  | 5% |
